# VaMiAnalyzer: An open source, python-based application for analysis of 3D *in vitro* vasculogenic mimicry assays

**DOI:** 10.1101/2025.05.13.653881

**Authors:** Stephen P.G. Moore, Xinyu Zhang, Olivia Chika Jonathan, Anqi Zou, Deborah Lang, Chao Zhang

## Abstract

**Background:** Vasculogenic mimicry (VM) is the phenomenon whereby non-vascular tumor cells develop vascular-like structures. VM is linked to more aggressive tumor phenotypes including higher rates of metastasis and invasion and is potentially resistant to anti-angiogenic cancer therapies. VM is investigated *in vitro* using 3D VM assays with microscopy images capturing the resulting VM structures. The standard method to quantify endpoint data is to count various structural features manually, which is time-consuming and open to bias. At present, no software solutions have been developed to specifically address the analysis and quantification of VM structures.

**Results:** To address this limitation, we developed an open source, python-based application, VaMiAnalyzer, allowing straightforward quantification of several VM structural features. The application follows a two-step approach that optionally corrects and enhances the raw input images and then analyzes and quantifies the VM features.

**Conclusions:** VaMiAnalyzer is stand-alone software that allows automated measurement of VM structural features from phase-contrast microscopy images. It produces results that are strongly consistent with manual counts but in a significantly shorter time, allowing speedy, non-biased analysis of VM from microscopy images.

## Background

Tumors require a constant blood supply to bring nutrients and oxygen, as well as remove waste products. This is achieved through the process of angiogenesis, the formation of new blood vessels from a pre-existing vascular network [1]. In contrast, vasculogenic mimicry (VM) is the phenomenon whereby existing, non-vascular tumor cells can form their own tube/loop structures allowing transportation of blood throughout the tumor [2, 3].

A common method to investigate VM formation is to perform *in vitro* assays, where cancer cells are grown on a “growth-factor reduced” 3D matrix of collagen or fibrin in the absence of nutrients and/or oxygen in 96-well cell culture plates [4]. The resulting VM structures are imaged by phase-contrast or fluorescence microscopy. However, no software solutions have been developed to specifically address the analysis and quantification of VM structures.

VM analysis studies often employ software adopted for angiogenesis assays, which share many characteristics with VM [5, 6]. To assess angiogenesis formation *in vitro*, several open source and commercial image analysis tools have been developed which can analyze and quantify vascular structures using several parameters such as number of branching points, tube number, and loop number [7]. However, VM results in less uniform vasculature structures compared to angiogenesis, due to VM originating from tumor cells that normally do not contribute to vasculature development. Furthermore, tumor cells (unlike endothelial cells) will continue to proliferate and form multilayered structures, leading to uneven focus in microscopy images. Due to these constraints, angiogenesis assay image analysis solutions do not work on VM images and fail to adequately and successfully quantitate VM-specific characteristics. This leads many VM researchers to manually count VM structures from microscope images, which is time consuming and subject to unintentional bias. To analyze VM quickly and accurately, new methodologies are needed. Here we describe an open source, python-based image analysis program that can quickly analyze and quantify VM structures using several parameters from microscope images without the potential addition of unintentional user bias.

## Implementation

The study employed a comprehensive two-step image analysis approach to quantify VM structures from microscope images (**Figure 1A**).

**Figure 1.**
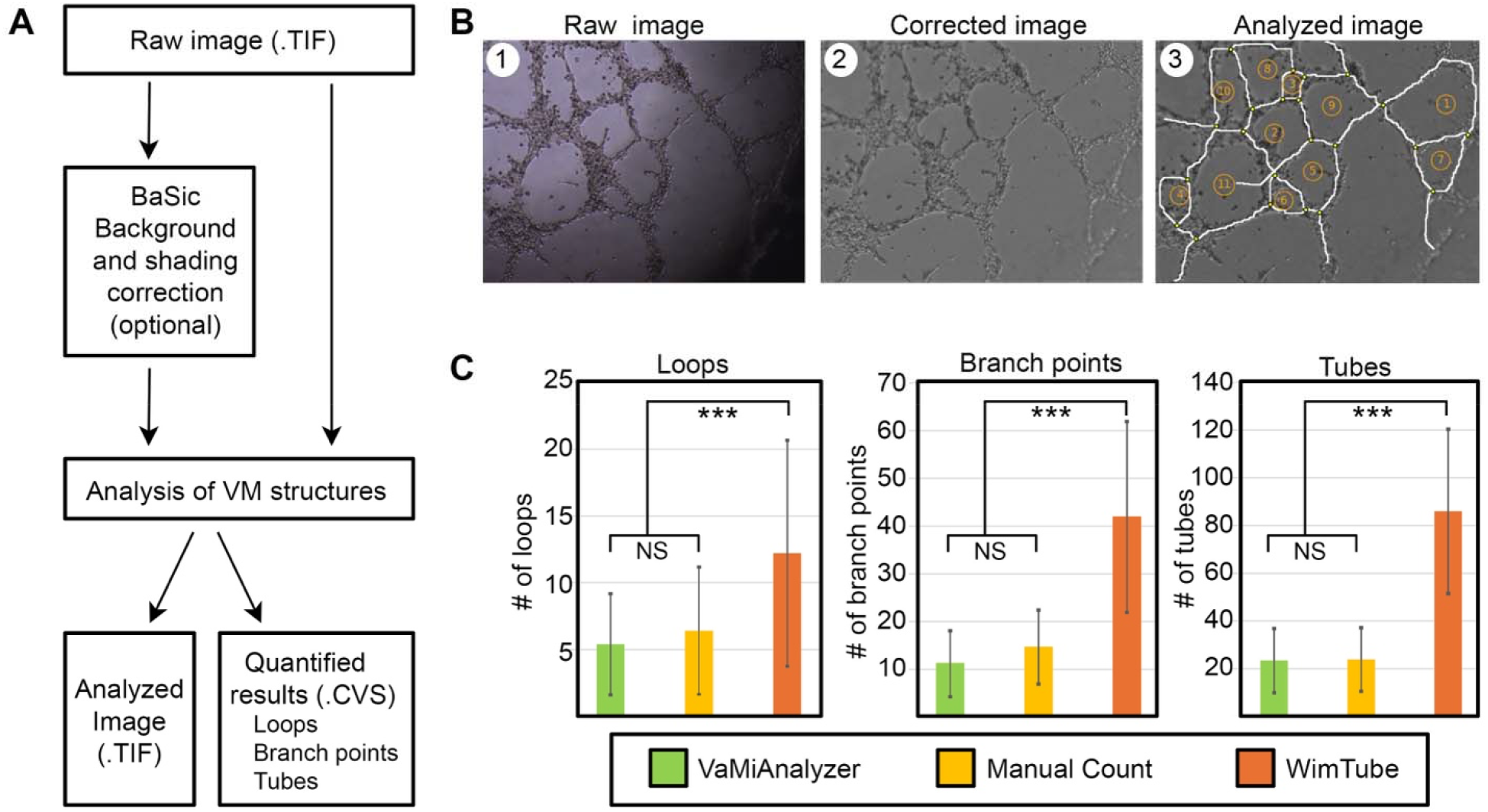
**(A)** Overview of VaMiAnalyzer two-step image analysis process. Raw images are analyzed for VM structures directly or after background and shading correction performed. **(B)** Representative images from each step of the analysis process. 1. Raw image input. 2. Corrected image after background and shading correction. 3. Analyzed image featuring tubes (white lines), branch points (yellow dots), and loop counts (orange numbers). **(C)** Comparison of VM analysis by VaMiAnalyzer, manual counting, and the WimTube tube formation AI image analysis program. Counts of Loops, Branch points, and Tubes did not differ significantly between the VaMiAnalyzer or manual counting of structures (p > 0.05). The WinTube program results were significantly different from the other two groups for all three structural features (p ≤ 0.05).

### Image Correction and Enhancement

The 3D structures formed in VM assays often result in images losing focus or having an uneven background, leading to biased quantification based on partial images (**Figure 1B**). This can affect analysis through both manual measurements and existing software designed for angiogenesis assay analysis. Image correction is necessary to ensure measurement quality and consistency. Each image underwent background and shading correction using the BaSiC algorithm [8], normalizing illumination and reducing artifacts. Variations in image brightness and shading were addressed to avoid affecting analyses. Mean and standard deviation values were adjusted for uniformity. Optional local contrast enhancement using Contrast Limited Adaptive Histogram Equalization was applied to improve the visibility of fine details and highlight subtle features crucial for accurate structure identification.

### VM Analysis and quantification

After correction, images were analyzed to identify and quantify tubular VM structures with the following steps:

- **Image Binarization and Structure Detection:** Images were binarized using a sliding window approach with the local variance method to optimize the detection of relevant structures. Parameters such as window size, step size, area threshold, and standard deviation threshold were carefully set to enhance detection accuracy. This method differentiated tubular structures from surrounding tissue based on pixel intensity variations, ensuring accurate identification and minimizing false positives. All parameters can be adjusted based on image resolution and structure size to optimize the outcome.
- **Graph Construction and Simplification:** Graphs constructed for quantitative analysis were based on the binarized VM structures. The binary images were first skeletonized to highlight tubular structures, reducing them to essential linear components while preserving their connectivity. This process transformed the images into thin representations of the original structures, maintaining the topology and critical pathways of the vascular formations. The Depth-First Search method was employed to traverse the skeleton, systematically exploring and mapping the entire network of vascular formations. During this traversal, marked key points were identified within the skeleton, specifically focusing on endpoints and junctions of the tubular structures. These points served as graphical nodes. Edges were defined as the segments connecting these nodes, capturing the pathways within the network. Nodes in close proximity were merged based on a predefined threshold, reducing graph complexity by eliminating redundant nodes. This process balanced the retention of important structural details with the need to minimize redundancy. Furthermore, shorter branches were pruned from the graph to eliminate noise and focus on significant vascular formations. This pruning enhanced the clarity and interpretability of the graph by removing extraneous elements.
- **Loops, Branch Points, and Tubular Structures Quantification:** The final step involves counting loops, branch points, and tubular structures to comprehensively quantify VM formation. Loops were identified using the NetworkX package [9], employing the Horton algorithm to find the minimum cycle basis by iteratively adding each edge to the shortest path between its endpoints. The resulting loops were filtered to retain only those with at least two branch points, ensuring that only significant loops contributing to the tubular network’s complexity were counted. Branch points were identified by examining the degree of each node in the graph, with nodes having more than two edges considered as branch points. By default, any two branch points closer than 50 pixels were merged to reduce redundancy and ensure accurate representation. Tubular structures were determined by counting edges in the pruned graph that exceeded a length threshold, set to 120 pixels and adjustable based on specific image characteristics. This ensured only significant tubular formations were considered. Each edge represented a tubular segment between two nodes, and short, insignificant segments were excluded from the count. The results were visualized and annotated to display identified structures and statistical data regarding loops, branch points, and tubes. Annotated images and statistical summaries provided a clear depiction of VM formation, complexity, and distribution within the samples.

## Results and Discussion

Human melanoma cell line MEL537 was cultured in DMEM (Gibco) with 10% FBS (Corning) without addition of antibiotics. Histological analysis was used to verify lack of mycoplasma contamination.

VM assays were performed as previously described (Francescone et al., 2011). Briefly, 50 μL of growth factor reduced Matrigel^®^ matrix (Corning) was added to the wells of a 96-well plate and incubated at 37°C for one hour to solidify. MEL537 tumor cells resuspended in DMEM without addition of FBS were added at 2 × 10^4^ cells per well and incubated at 37°C in 5% CO2 for 24 hours. Four images were captured from random positions in each well using an Olympus QColor 3 camera mounted to an Olympus CKX41 microscope system through a 10x objective.

VM assays are an *in vitro* method used to measure the formation of VM by aggressive cancer cells, and to test the impact that inhibition of target gene/s, or potential therapeutic compounds have on this phenomenon. VM analysis is often implemented by manual counting of structures due to a lack of suitable analysis tools. We developed VaMiAnalyzer to enable automated, unbiased quantification of VM structures from 3D VM assay images (**Figure 1A**). To test the functionality of VaMiAnalyzer we performed VM assays using MEL537 melanoma cells in standard 96-well plate based Matrigel^®^ VM assays; images were taken from random points within several wells. Since VM assays are typically performed in 96-well plates and multiple images are taken per well [1], corner shadowing from the sides of the wells can occur (**Figure 1B)**. To counter this, image correction and enhancement of the raw input image by VaMiAnalyzer can be performed to produce uniform shading, brightness, and contrast across the entire image (**Figure 1B**). However, if no shading is evident in images the correction and enhancement feature may be omitted. VM structural analysis by VaMiAnalyzer measures and quantifies three key features: the number of loop structures, branching points, and tube structures (**Figure 1B**). Quantified results are outputted as csv documents to the specified output folder with count data for each structural type per image, thereby allowing for easy averaging and statistical analysis.

Comparison of average count data for each structural type between VaMiAnalyzer, manual counting, and the commercial WimTube tube formation analysis program (https://www.wimasis.com) was performed on 15 representative VM images. Image analysis of loops, branch points, and tube numbers by VaMiAnalyzer was comparable to manual counting, while the commercial program overestimates all three structure types when compared to the other two methods. One-way ANOVA and Tukey post-hoc analysis found no significant difference in counts of any of the three VM structural features analyzed by VaMiAnalyzer or manual counting, while WimTube counts were significantly different from either VaMiAnalyzer or manual counts (**Figure 1C**). These results demonstrate that VaMiAnalyzer is an appropriate method for VM analysis with equivalent results to manual analysis. Additionally, adoption of VaMiAnalyzer can save a substantial amount of time. Manual counting of the 15 VM images took an average of 11.8 minutes per image, whereas VaMiAnalyzer averaged 43.5 seconds per image (16 second average for correction, 27.5 second average for analysis) a reduction of over 11 minutes per image, or 2.8 hours for our total 15 image dataset. In short, VaMiAnalyzer provides a robust, easy to use, and automated solution for analysis of 3D VM assays from microscopy images and eliminates the potential for unintentional biasing of results. VaMiAnalyzer was developed using python version 3.x. It depends on functions from the python packages *OpenCV, NumPy, BaSiC, Matplotlib, Scikit-image, NetworkX*, and *TQDM*. The software is platform-independent and easy to install. The source code, user manual, installation instructions, testing data, and tutorial are freely available at GitHub.

## Conclusions

VaMiAnalyzer is a stand-alone, python-based application designed to quantify several structural features that form during VM development in specialized 3D assays. Due to the nonuniformity of VM structural development by tumor cells, pre-existing software developed for similar but more uniform angiogenesis assays does not accurately quantify VM structures, with researchers often relying on manual counting instead. This easy-to-use application allows automated measurement of VM structures from phase-contrast microscopy images, producing results comparable to manual counting in a significantly shorter time while eliminating the potential biasing of results.

## Declarations

### Ethics approval and consent to participate

Not applicable

### Consent for publication

Not applicable

### Availability and Requiements

Project name: VaMiAnalyzer

Project home page: https://github.com/CZCBLab/VaMiAnalyzer

Operating system(s): Platform independent

Programming language: python

Other requirements: python 3.0 or higher

License: GNU GPL v3.

Any restrictions to use by non-academics: licence needed

## Data Availability

The images and results generated and analysed during the current study are available at: https://github.com/CZCBLab/VaMiAnalyzer

## Authors’ contributions

S.PG.M., D.L., and C.Z. conceived and designed the project. X.Z. and A.Z. developed the image processing method and implemented the software. S.PG.M. and O.C.J. generated the images and conducted the manual measurements. S.PG.M., X.Z., D.L., and C.Z. wrote the manuscript. All authors reviewed the final manuscript.

## Acknowledgements

Not applicable

## References

1. Tonini T, Rossi F, Claudio PP: Molecular basis of angiogenesis and cancer. Oncogene 2003, 22(42):6549–6556.

2. Hendrix MJ, Seftor EA, Seftor RE, Chao JT, Chien DS, Chu YW: Tumor cell vascular mimicry: Novel targeting opportunity in melanoma. Pharmacol Ther 2016, 159:83–92.

3. Folberg R, Hendrix MJ, Maniotis AJ: Vasculogenic mimicry and tumor angiogenesis. Am J Pathol 2000, 156(2):361–381.

4. Francescone R, Vendramini-Costa DB: In Vitro Models to Study Angiogenesis and Vasculature. Methods Mol Biol 2022, 2514:15–28.

5. Kubota Y, Kleinman HK, Martin GR, Lawley TJ: Role of laminin and basement membrane in the morphological differentiation of human endothelial cells into capillary-like structures. J Cell Biol 1988, 107(4):1589–1598.

6. Arnaoutova I, Kleinman HK: In vitro angiogenesis: endothelial cell tube formation on gelled basement membrane extract. Nat Protoc 2010, 5(4):628–635.

7. Pereira M, Pinto J, Arteaga B, Guerra A, Jorge RN, Monteiro FJ, Salgado CL: A Comprehensive Look at In Vitro Angiogenesis Image Analysis Software. Int J Mol Sci 2023, 24(24).

8. Peng T, Thorn K, Schroeder T, Wang L, Theis FJ, Marr C, Navab N: A BaSiC tool for background and shading correction of optical microscopy images. Nat Commun 2017, 8:14836.

9. Hagberg A, Swart PJ, Schult DA: Exploring network structure, dynamics, and function using NetworkX. In: Conference: SCIPY 08; August 21, 2008; Pasadena, Pasadena, CA (United States), 21 Aug 2008; United States. AC52-06NA25396 2024-02-01: 2008: Medium: ED.

